# β-hydroxybutyrate modulates enteric pathogen susceptibility through regulation of commensal bacteria and intestinal Th17 responses

**DOI:** 10.64898/2026.03.20.713262

**Authors:** Wenxuan Dong, Chi Yan, Bethany Korwin-Mihavics, Kate Stack, Gillian Hughes, Aidan Schmidt, Kevin Schwartz, Gustavo Caballero-Flores, Margaret Alexander

**Affiliations:** Department of Medical Microbiology & Immunology, University of Wisconsin-Madison, Madison, Wisconsin, 53706, USA

## Abstract

T helper 17 (Th17) cells are a critical T lymphocyte subset involved in mucosal immunity and host defense against enteric pathogens. Although ketogenic diets (KD) and the major ketone body β-hydroxybutyrate (BHB) reshape gut microbiota and suppress Th17 responses under defined diet conditions, it remains unclear whether elevation of BHB alone, independent of dietary macronutrient composition and systematic metabolic shift, is sufficient to remodel Th17-inducing commensals and alter host susceptibility to enteric infection. Here, we used 1,3-butanediol (BD), a precursor metabolized to BHB independently of KD, to elevate systemic BHB levels in mice. BD treatment significantly reduced the frequency of ileal Th17 cells, as assessed by flow cytometry for Th17 markers IL-17A and RORγt. 16S rRNA gene sequencing revealed that BD altered gut microbial community structure, as indicated by beta-diversity analysis based on Bray-Curtis dissimilarity, and reduced Shannon diversity and evenness. Linear discriminant analysis effect size identified segmented filamentous bacteria (SFB) as significantly decreased in the ileum following BD treatment, and SFB abundance positively correlated with Th17 markers. Microbiota transplantation demonstrated that BD-shaped microbiota was sufficient to suppress Th17 responses in recipient mice, accompanied by reduced SFB abundance. In a *Citrobacter rodentium* infection model, BD treatment was associated with increased pathogen burden, and fecal *C. rodentium* levels were negatively correlated with SFB abundance. Together, these results indicate that BD-induced elevation of BHB reshapes commensal microbiota, including decreasing SFB levels, resulting in dampened Th17 responses and increased susceptibility to enteric infection.

**IMPORTANCE:** Diet is a key determinant of gut microbial composition and mucosal immune function, yet the microbial mechanisms linking how diet-mediated changes to metabolism regulate immune responses remain incompletely understood. Th17 cells play central roles in both protective mucosal immunity and inflammatory pathology, making them a critical target of immunometabolic regulation. In this study, we show that β-hydroxybutyrate (BHB), generated independently of diet, suppresses intestinal Th17 responses by reshaping the gut microbiota, reducing SFB levels, a potent Th17-inducing murine commensal. We further demonstrate that BHB-associated microbiota changes are linked to increased susceptibility to enteric infection. This work provides a mechanistic framework illustrating how metabolic state can influence host immunity through selective effects on commensal microbes. These findings inform future studies of microbiota-mediated immune regulation.

## OBSERVATION

Dietary intervention has emerged as a promising strategy for modulating immune responses in inflammatory diseases. T helper 17 (Th17) cells play context-dependent roles in immunity, contributing to host defense against enteric pathogens while also driving pathogenic inflammation in autoimmune diseases (1–3). As a result, precise regulation of Th17 responses has been proposed as a means to balance protective immunity and inflammatory pathology. High-fat low-carbohydrate ketogenic diets (KDs) that shift host metabolism to ketogenesis can alleviate autoimmune phenotypes through suppression of Th17 cells, an effect that has been associated with alterations in the gut microbiota (4–6). However, KDs profoundly change macronutrient composition and host metabolism, making it difficult to disentangle the immunological effects of ketogenesis from those of dietary fat and carbohydrate restriction. β-Hydroxybutyrate (BHB), the major circulating ketone body produced during ketogenesis, functions as a signaling metabolite with roles in gene regulation and immune modulation, including effects on inflammasome activation and innate lymphoid cells (7–9). KDs and BHB also enhance antiviral immunity by modulating T cell metabolism and function, improving disease outcomes and survival during infections (10, 11). However, whether BHB alone is sufficient to regulate intestinal Th17 responses independently of diet and the impact on enteric infection susceptibility remains unclear. To address this question, we used 1,3-butanediol (BD), a metabolic precursor that elevates systemic BHB levels without altering dietary macronutrient composition, to isolate the effects of BHB from those of diet (12). Using a combination of flow cytometry, 16S rRNA gene sequencing, microbiota transplantation, and *Citrobacter rodentium* infection models, we examined how BHB influences Th17 responses and host susceptibility to enteric infection.

### Isolated elevation of BHB suppresses intestinal Th17 responses and reshapes the gut microbiota

To test whether BHB suppresses Th17 cells independently of diet, mice were supplemented with 20% BD in drinking water to elevate systemic BHB levels (12–14) (Fig. 1A). Compared to control mice (CON), BD-treated mice exhibited a significant reduction in ileal Th17 cell frequency, as defined by CD4^+^IL-17A^+^ and CD4^+^RORγt^+^ populations among total live CD3^+^ cells (Fig. 1B; Fig. S1). These results indicate that elevation of BHB alone is sufficient to suppress intestinal Th17 responses in the absence of other dietary alterations.

**Figure 1.**
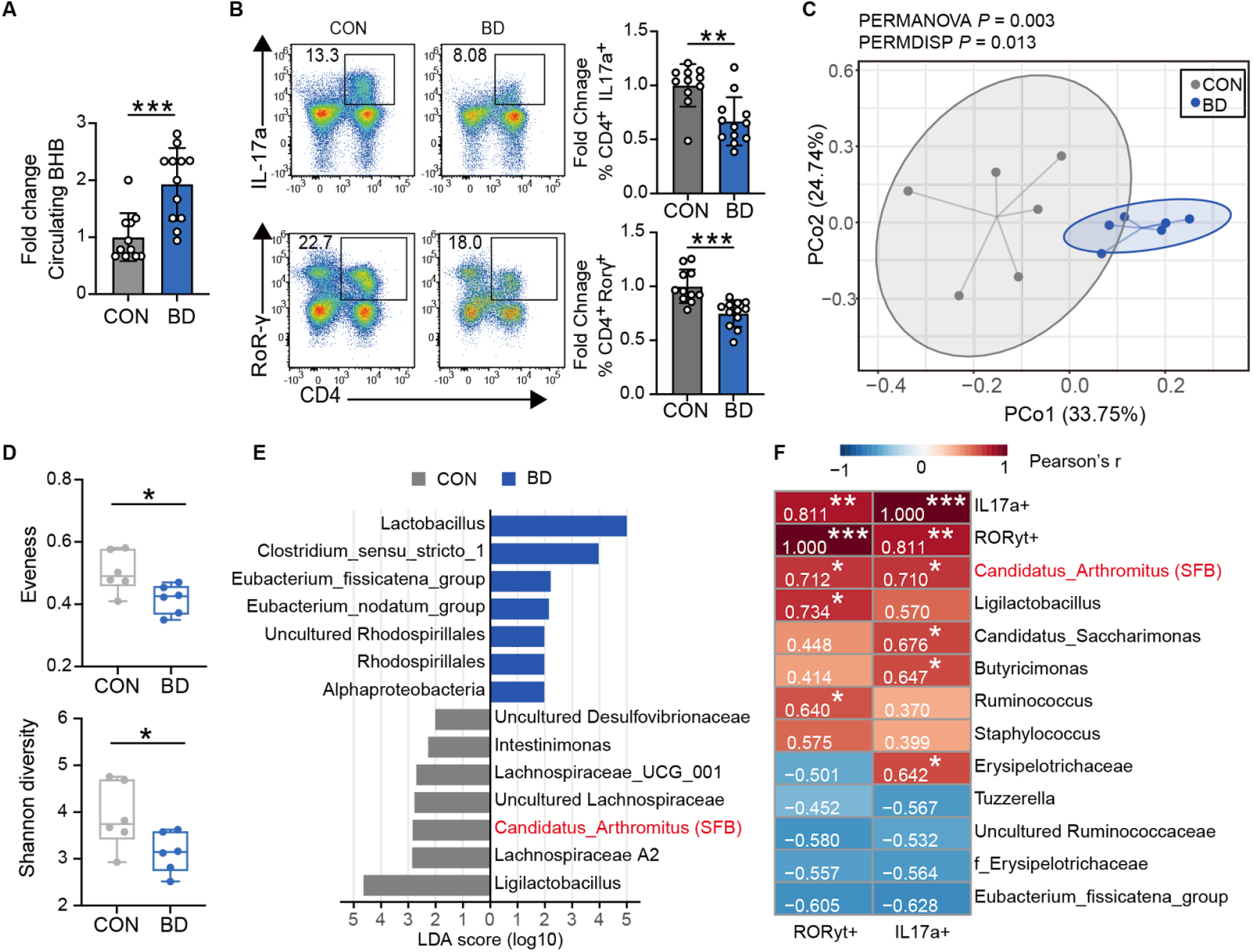
Elevated BHB decreases ileal Th17 levels and reshapes the microbiota. (**A**) Fold-change in circulating β-hydroxybutyrate (BHB) levels in control (CON) and 1,3-butanediol (BD)-treated mice. Barplot shows mean ± standard deviation (SD). BD significantly increased circulating BHB compared to CON. Statistical analysis: unpaired two-tailed Welch’s t test. (**B**) Flow cytometry analysis of ileal lamina propria lymphocytes. Left, representative flow plots showing CD4^+^IL-17A^+^ and CD4^+^RORγt^+^ cells gated on live CD3^+^ populations. Right, quantification of Th17 frequency normalized to CON. Barplots show mean ± SD. Statistical analysis: unpaired two-tailed Welch’s t test. (**C**) Principal coordinates analysis (PCoA) based on Bray-Curtis dissimilarity showing differences in β-diversity between CON and BD groups. PERMANOVA and PERMDISP *p*-values are indicated. (**D**) Alpha diversity analysis of ileal microbiota. BD reduced community evenness (top) and Shannon diversity (bottom). Boxplots show the median, interquartile range (IQR), and whiskers represent the minimum and maximum values. Statistical analysis: Mann-Whitney U test. (**E**) Differentially abundant genera identified by linear discriminant analysis effect size (LEfSe). Bars represent LDA scores (log10 scale), and colors indicate taxa enriched in each group. (**F**) Pearson correlation heatmap between selected bacterial genera (|correlation coefficient| > 0.5 and P < 0.1 for correlation with either RORγt^+^ or IL-17A^+^ CD4^+^ cells) and Th17 markers (IL-17A^+^ and RORγt^+^ CD4^+^ cells). Segmented filamentous bacteria (SFB; annotated as *Candidatus Arthromitus*) is highlighted in red. Numbers represent Pearson’s correlation coefficients. \**p* < 0.05, \*\**p* < 0.01, \*\*\**p* < 0.001.

Given the well-established role of the gut microbiota in Th17 cell response, we next examined whether BD-mediated BHB production altered the intestinal microbial community. 16S rRNA gene sequencing revealed that BD significantly changed ileal microbiota composition compared to CON (Fig. 1C; Fig. S2). PERMANOVA revealed a significant difference in community composition between groups, and PERMDISP indicated reduced beta-dispersion in BD group, suggesting an increased microbiome homogeneity. BD was also associated with reduced alpha diversity, including lower Shannon diversity and evenness (Fig. 1D). Previous studies have shown reduced Th17 cells in germ-free and antibiotic-treated mice and specific members of the microbiota are known to elevate Th17 levels (15, 16), raising the possibility that BD-mediated suppression of Th17 responses may be mediated by microbiota alterations. Together, these results suggest that BD reshapes the gut microbiota, resulting in a less diverse and more compositionally homogeneous community associated with reduced Th17 responses.

### BHB-associated Th17 suppression coincides with reduced segmented filamentous bacteria (SFB)

Differential abundance analysis using linear discriminant analysis effect size (LEfSe) identified segmented filamentous bacteria (SFB, annotated as “*Candidatus arthromitus”*) as one of the most significantly reduced taxa in BD-treated mice (Fig. 1E). SFB are well-characterized commensals that potently induce Th17 activation through adherence to small intestinal epithelial cells and subsequent antigen presentation by dendritic cells (17, 18). Consistent with this role, correlation analysis demonstrated that SFB was the only taxa that positively correlated with both Th17 markers assessed in the ileum (Fig. 1F). These findings suggest that BD-induced BHB suppresses Th17 responses in part through reduction of SFB levels.

To determine whether the altered microbiota under BD was sufficient to recapitulate Th17 suppression, we performed microbiota transplantation experiments. Recipient mice colonized with microbiota from BD donors had significantly lower ileal Th17 frequency compared to recipients of CON microbiota (Fig. 2A). Quantitative PCR analysis revealed that absolute SFB abundance was significantly lower in recipients of BD-shaped microbiota (Fig. 2B). As observed in donor mice, SFB levels in recipients positively correlated with ileal Th17 frequency (Fig. 2C), indicating that the BD-altered microbiota is sufficient to transmit both reduced SFB levels and suppressed Th17 responses. These results support a microbiota-dependent mechanism by which BHB modulates intestinal Th17 immunity, with reduction of SFB representing one prominent component. This observation extends prior work demonstrating that KD or BHB ketone ester supplementation can influence Th17 by increasing *Lactobacillus murinus* or reducing levels of *Bifidobacterium* under high-fat diet conditions (4, 5). Together, these studies suggest that ketogenesis may reduce Th17 responses through multiple microbiota-dependent pathways, the relative contribution of which may depend on the ecological configuration of the host microbiome. Our results add an additional layer of complexity by identifying SFB reduction as a distinct microbiota axis through which BHB elevation can influence Th17 levels in the intestine.

**Figure 2.**
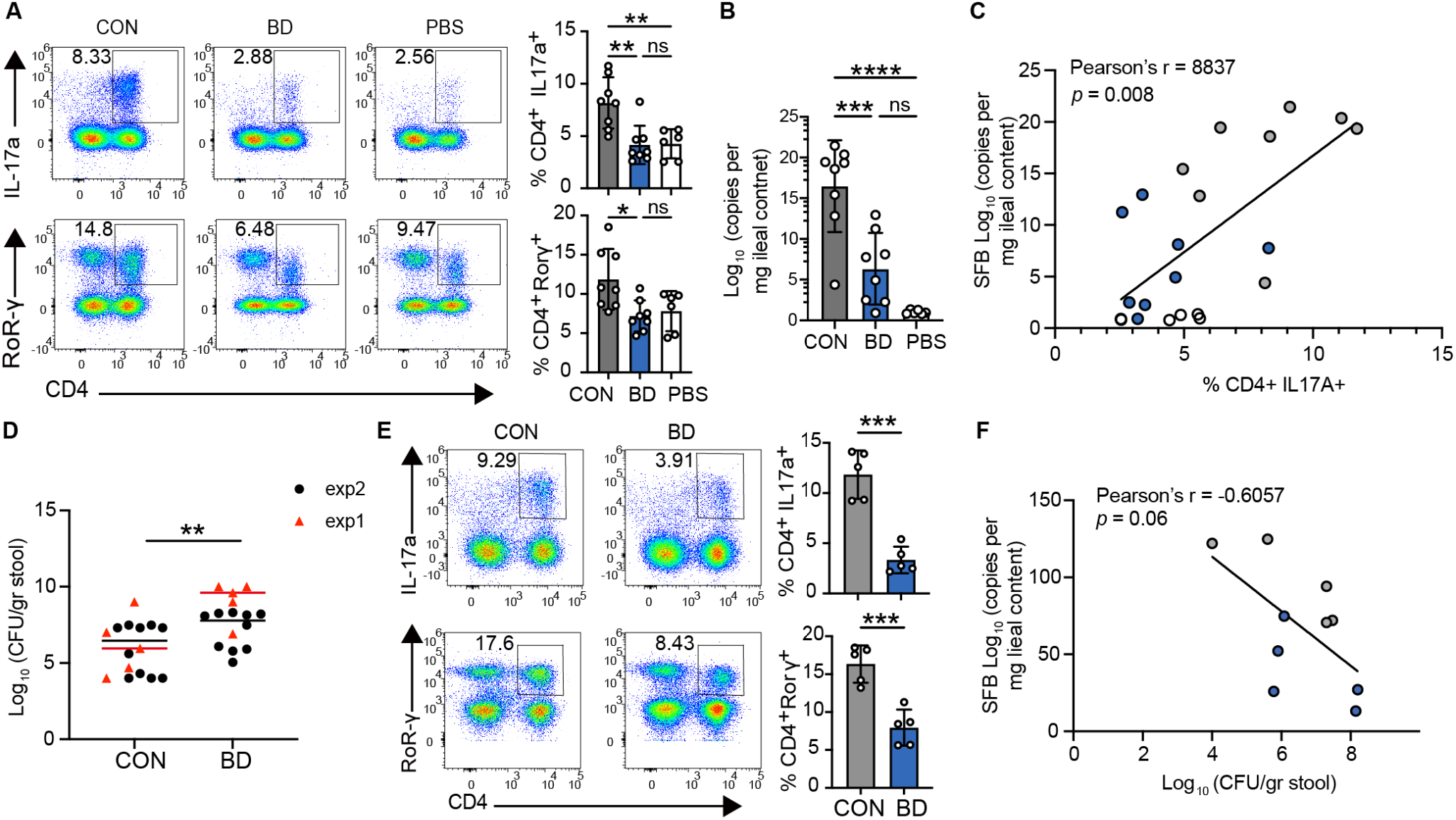
BHB alters *Citrobacter rodentium* susceptibility in association with reduced SFB abundance and Th17 responses. (**A**) Flow cytometry analysis of ileal lamina propria lymphocytes following microbiota transplantation or a PBS control. Left, representative flow plots showing CD4^+^IL-17A^+^ and CD4^+^RORγt^+^ cells gated on live CD3^+^ populations. Right, quantification of Th17 frequencies. Barplots show mean ± SD. Statistical analysis: one-way ANOVA. (**B**) Quantitative PCR analysis of segmented filamentous bacteria (SFB) absolute abundance in recipient mice. Barplots show mean ± SD. Recipients of BD-shaped microbiota exhibited reduced SFB levels. Statistical analysis: one-way ANOVA. (**C**) Pearson correlation analysis showing a negative correlation between SFB abundance and ileal Th17 frequency. Colors indicate recipient groups. (**D**) Fecal *C. rodentium* burden 4 days post-infection in CON and BD-treated mice. Mice were pre-treated with BD water 7 days prior to infection. Colors indicate two independent experimental repeats. Horizontal bars indicate the medians for each independent experiment. Statistical analysis: unpaired two-tailed Welch’s t test. (**E**) Flow cytometry quantification of ileal Th17 frequency on day 4 post-infection. Barplots show mean ± SD. Statistical analysis: unpaired two-tailed Welch’s t test. Left, representative flow plots showing CD4^+^IL-17A^+^ and CD4^+^RORγt^+^ cells gated on live CD3^+^ populations. Right, quantification of Th17 frequencies. Statistical analysis: unpaired two-tailed Welch’s t test. (**F**) Pearson correlation between SFB abundance and fecal *C. rodentium* burden. \**p* < 0.05, \*\**p* < 0.01, \*\*\**p* < 0.001.

### BHB alters *C. rodentium* susceptibility in association with reduced SFB and Th17 responses

Th17 cells and their effector cytokines, including IL-17A, have been implicated in host defense against enteric pathogens such as *Salmonella* and *C. rodentium* (19). SFB colonization has been shown to confer protection against *C. rodentium* infection, in part by enhancing Th17 responses (17). To examine how BD-induced alterations in microbiota and Th17 responses relate to host susceptibility to infection, we challenged mice that had been pre-treated with BD or CON water for 7 days with *C. rodentium* (20, 21). BD pre-treated mice exhibited significantly increased pathogen burden on day 4 post infection and increased mortality compared to CON group (Fig. 2D, Fig. S3). This corresponded with reduced intestinal Th17 levels in BD treated mice during infection (Fig. 2E). Further, correlation analysis revealed a negative correlation between fecal *C. rodentium* load and intestinal SFB abundance (Fig. 2F). Consistent with observations in uninfected mice, BD treatment also reduced SFB abundance following *C. rodentium* infection, and SFB levels remained positively correlated with Th17 markers in the ileum (Fig. S4). Together, these findings suggest that BD-mediated SFB reduction dampens Th17 responses and increases susceptibility during enteric infection.

Notably, the increased pathogen burden observed under BD coincided with reduced SFB and suppressed Th17 responses, although prior studies have demonstrated that SFB-mediated protection against *C. rodentium* during early infection can occur independently of CD4^+^ T cells (22). Thus, our findings are consistent with a model in which BD-induced BHB disrupts SFB-associated colonization resistance and immune priming, leading to increased susceptibility to enteric infection, while concurrently suppressing Th17 responses. These results extend prior work demonstrating that ketogenic metabolism can modulate host-pathogen interactions in a tissue-specific manner. For example, KD or BD supplementation has been shown to expand protective γδ T cells in the lung and enhance barrier function, thereby improving resistance to viral infection (10). In contrast, our data reveal in the context of an enteric pathogen, elevated BHB results in increased pathogen burden, suggesting an opposing tissue-specific response.

In summary, our results demonstrate that elevation of BHB via BD is sufficient to suppress intestinal Th17 responses independent of diet composition. This effect is associated with remodeling of the gut microbiota, particularly a reduction of the Th17-inducing commensal SFB, and is transferable through microbiota transplantation. In the context of *C. rodentium* infection, BD-induced microbiota changes are linked to increased pathogen burden, highlighting a potential trade-off between suppression of inflammatory Th17 responses and maintenance of effective colonization resistance. Together, these findings define a diet-independent ketogenesis–microbiota–immune axis that shapes intestinal Th17 immunity and host-pathogen interactions in mice.

## DATA AVAILABILITY

All 16S rRNA gene sequencing data are available at the NCBI Sequence Read Archive under BioProject No. PRJNA1429443. This paper does not report original code.

## SUPPLEMENTAL MATERIAL

Supplemental Figures S1-S4

Materials and Methods

## ACKNOWLEDGMENTS

This research was supported by NIH grant R00AI159227, Shaw Early Research Career Award (Greater Milwaukee Foundation), WARF/OVCR and UW-Madison Start-up funding to M.A. Work by G. C.-F. and A.S. was supported by NIH grant R00AI159620 and UW-Madison Start-up funds to G. C.-F.

We thank the Gumperz Lab in the Department of Medical Microbiology & Immunology at UW-Madison for their assistance and technical support with flow cytometry analyses. The authors thank the University of Wisconsin Carbone Cancer Center Flow Cytometry Laboratory, supported by P30 CA014520, for use of its facilities and services.

