## Supplemental Figures 1-4 for "β-hydroxybutyrate modulates enteric pathogen susceptibility through regulation of commensal bacteria and intestinal Th17 responses"

### Supplemental Information (SI)

#### Supplemental Figures

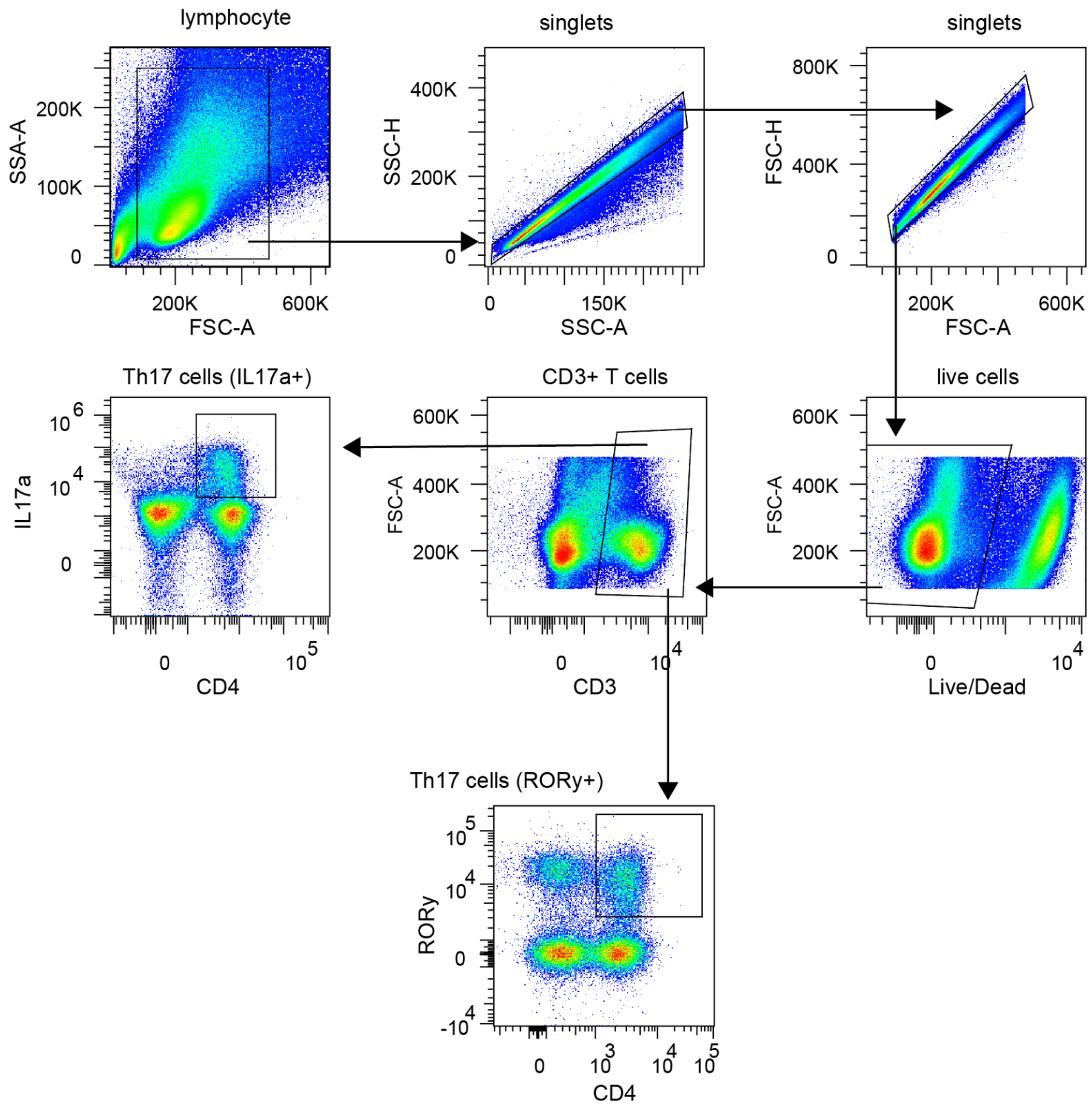


**Fig. S1.** Gating strategy for Th17 cell populations. Cells were gated on lymphocytes, single cells, single cells, live cells, CD3+ cells, then CD4+ IL-17A+ cells or CD4+ RORγt+.

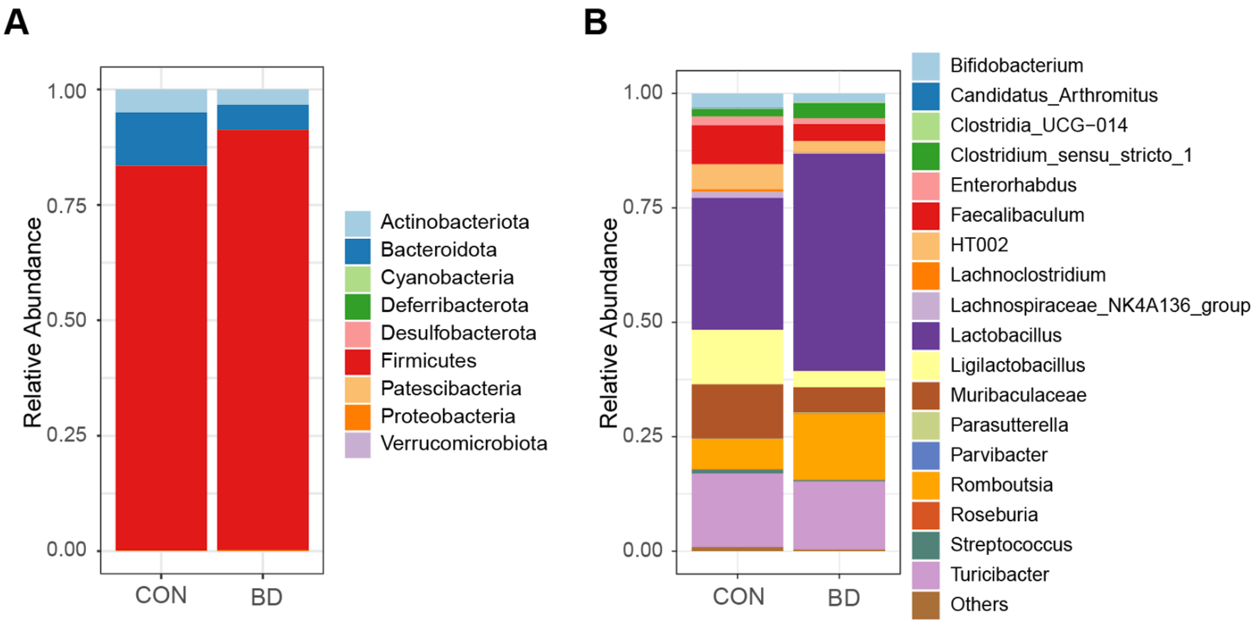


**Fig. S2.** Stacked bar plots depicting the relative abundance of bacterial taxa in control (CON) and 1,3-butanediol (BD)-treated mice at the phylum level (**A**) and genus level (**B**).

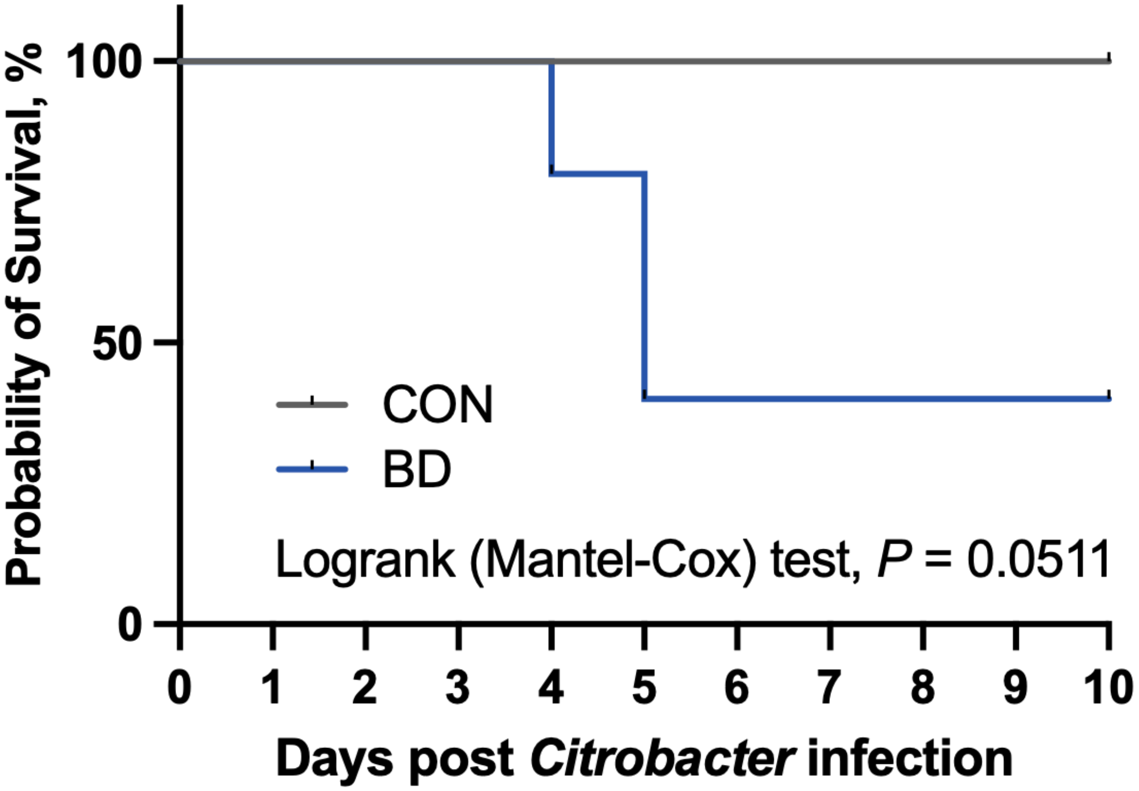


**Fig. S3**. BD pretreatment decreased the probability of survival post *C. rodentium* infection. Data are from experiment 1 (n=5). Statistics: Logrank (Mantel-Cox) test.

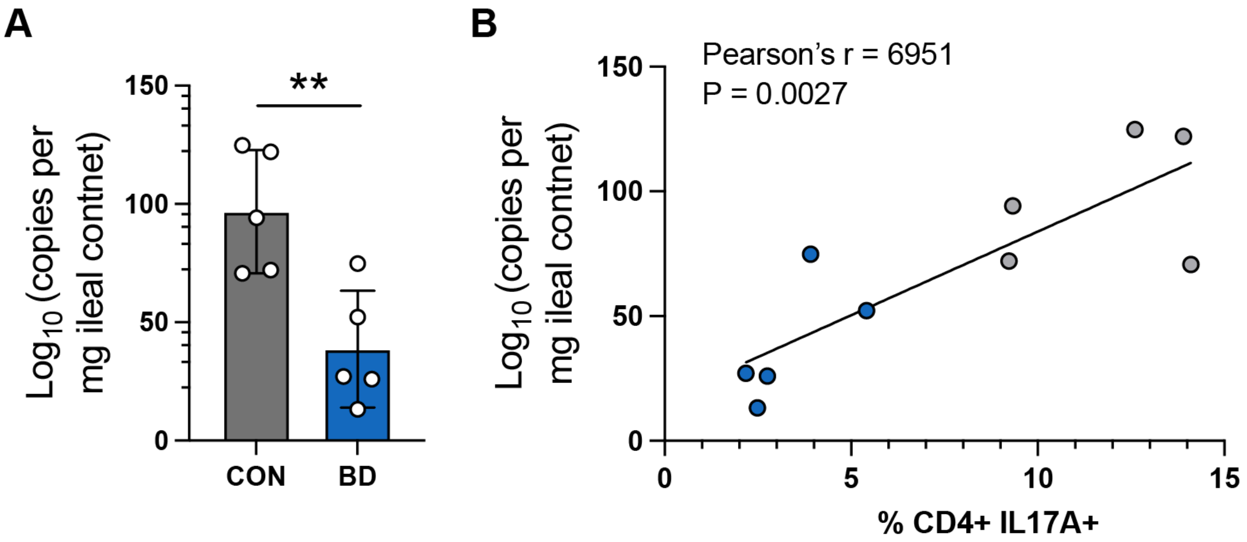


**Fig. S4.** BD reduces ileal SFB abundance and SFB positively correlates with Th17 responses during *C. rodentium* infection. (**A**) Quantitative PCR analysis showing reduced segmented filamentous bacteria (SFB) abundance in the ileal contents of BD-treated mice 4 days after *C. rodentium* infection. (**B**) Pearson correlation analysis demonstrating a positive correlation between ileal SFB abundance and the frequency of $CD4^{+}IL\text{-}17A^{+}$ cells gated on live $CD3^{+}$ on day 4 post-infection.
