## Supplemental Materials - Methods for "β-hydroxybutyrate modulates enteric pathogen susceptibility through regulation of commensal bacteria and intestinal Th17 responses"

### Supplemental Information (SI)

#### Materials and Methods

##### Mice

All animal experiments were approved by the University of Wisconsin-Madison (UW-Madison) Institutional Animal Care and Use Committee (IACUC protocol M006785). C57BL/6 mice were bred at the UW-Madison Biomedical Research Model Services (BRMS) facility or purchased from Taconic Biosciences (C57BL/6 Tac). All mice were housed in the UW-Madison Microbial Sciences Building animal facility under specific pathogen-free (SPF) conditions. Mice were maintained on a 12-hour light/12-hour dark cycle with *ad libitum* access to standard chow (Teklad Irradiated Global 19% Protein Extruded Rodent Diet, #2919; Inotiv) and acidified water. Mice used in experiments were 7-9 weeks of age and were assigned to experimental groups to achieve similar age distribution. No animals had been subjected to prior experimental procedures.

##### 1,3-butanediol mouse experiment (Figure 1)

C57BL/6 Tac mice (equal numbers of males and females, 7-9 weeks old) were provided either standard acidified drinking water (CON) or drinking water supplemented with 20% (vol/vol) 1,3-butanediol (BD, Sigma B84785) for 14 days. Male and female mice were studied in separate experimental cohorts conducted at different time points; all other experimental conditions were identical between sexes. Mice were euthanized on day 14 post treatment for tissue collection. Ileal lamina propria lymphocytes were isolated for quantification of Th17 cells by flow cytometry. Ileal luminal contents were collected for 16S rRNA gene sequencing. Circulating BHB levels were measured using the Precision Xtra blood glucose and ketone monitoring system (Abbott; cat# 98814-65) with blood b-Ketone test strips (Abbott; cat# ART07249) on day 14. Tail blood was applied directly to test strips according to the manufacturer's instructions.

##### Microbiota transplantation experiment

Donor mice (C57BL/6 Tac) were maintained on either control drinking water (CON) or water supplemented with 20% (vol/vol) BD for 14 days prior to intestinal content collection. Donor mice (n = 8 per group; 4 males and 4 females) were cohoused by treatment group and sex. On days 0 and 14, half of the donor mice were euthanized for collection of ileal and cecal luminal contents.

Donor materials were prepared as previously described (1). Briefly, ileal and cecal contents were resuspended at a ratio of 1:10 (g/mL) in sterile, pre-reduced 1X phosphate-buffered saline (PBS) within an anaerobic Coy chamber. Samples were vortexed for 1 minute, allowed to settle for 2 minutes, and the supernatants were collected for transplantation. Recipient mice received 100 µL of ileal suspension by oral gavage in the morning and 100 µL of cecal suspension in the afternoon on the same day. In addition, soiled bedding from donor cages was transferred to recipient cages.

Transplantation procedure was conducted as previously described with modifications (2). Recipient C57BL/6 mice (bred at UW-Madison BRMS) were pretreated with antibiotics in drinking water for 7 days prior to microbiota transplantation. Antibiotics cocktail (AVNM) consists of ampicillin (1 g/L), vancomycin (0.5 g/L), neomycin (1 g/L), metronidazole (1 g/L) and sucrose (10 g/L). Antibiotic-containing water was prepared in sterile drinking water and filter-sterilized (0.2 µm). Following a 2-day antibiotic washout period, microbiota transplantation (MT) was performed as described above.

Recipient groups included CON microbiota recipients (n = 8), BD microbiota recipients (n = 8), and sterile PBS gavage controls (n = 6). All recipient mice were euthanized 28 days after the first gavage. Ileal lamina propria lymphocytes were isolated for Th17 quantification, and ileal contents were collected for quantitative PCR (qPCR) analysis.

##### ***Citrobacter rodentium*** **infection experiments**

*Citrobacter rodentium* DBS120 *(C. rodentium)* infection experiments were performed according to previously described (3). C57BL/6 mice were maintained on either control drinking water (CON) or water supplemented with 20% (vol/vol) BD for 7 days prior to *C. rodentium* infection. For infection experiments, *C. rodentium* was grown in Luria-Bertani (LB) broth at 37°C with shaking to an optical density at 600 nm (OD600) of approximately 1.0. Cultures were centrifuged, washed once with sterile PBS, and resuspended in PBS to the indicated concentrations (~5 × $10^9$ CFU/mL). Mice were orally gavaged with 200 uL bacterial suspension per animal. Fecal pellets were collected at designated time points post-infection, homogenized in PBS, serially diluted, and plated on LB agar containing kanamycin to determine bacterial burdens.

In experiment 1, all mice were obtained from Taconic Biosciences (n = 5 per group). In experiment 2, mice were bred at UW-Madison BRMS (n = 10 per group) and exposed to soiled bedding from Taconic mice for 10 days prior to BD supplementation and infection to standardize microbiota composition. In experiment 2, a subset of mice was euthanized on day 4 post-infection for isolation of ileal lamina propria lymphocytes and collection of ileal luminal contents for Th17 analysis and qPCR of segmented filamentous bacteria (SFB), as described below.

##### Lamina propria lymphocytes isolation

Lamina propria lymphocytes were isolated from ileum with minor modifications of previously described methods (4). Briefly, Peyer's patches were removed and the lower two-thirds of the small intestine was opened longitudinally and cleared luminal contents. Tissues were maintained in complete RPMI 1640 supplemented with 10% fetal bovine serum (FBS, Gibco A5256801), penicillin and streptomycin (100 U/mL), GlutaMax, sodium pyruvate, and nonessential amino acids.

To remove epithelial cells, intestinal tissue was incubated in calcium- and magnesium-free Hank's balanced salt solution (HBSS) containing 5 mM EDTA and 1 mM dithiothreitol (DTT) for 45 minutes at 37°C with shaking (200 rpm). Following filtration through a 100-μm cell strainer, tissues were further digested in HBSS supplemented with 5% FBS, Dispase (1 U/mL), collagenase VIII (0.5 mg/mL), and DNase I (20 μg/mL) for 35 minutes at 37°C with agitation. The resulting cell suspension was passed through a 40-μm cell strainer and washed in PBS. Mononuclear cells were enriched using a 40%/80% (vol/vol) Percoll density gradient centrifuged at 2,000 rpm for 20 minutes without brake. Cells collected at the interface were washed with PBS and processed for flow cytometric analysis as described below.

##### Flow cytometry

For surface staining, cells were incubated in a staining buffer (HBSS supplemented with 10 mM HEPES, 2 mM EDTA, and 0.5% FBS) for 20 minutes at 4°C. The following antibodies were used for surface staining: anti-CD3 (clone 17A2), and anti-CD4 (clone GK1.5). Cell viability was assessed using LIVE/DEAD Fixable Dead Cell Stain (Thermo Fisher Scientific).

For intracellular cytokine detection, cells were stimulated with Cell Stimulation Cocktail (eBioscience™, Invitrogen, cat# 00-4970-93), and brefeldin A (Golgi Plug; BD Biosciences, cat# 555029) for 4 hours at 37°C. Cells were then surface stained, fixed, and permeabilized using Invitrogen fixation/permeabilization buffer (cat# 00-5523-00) according to the manufacturer's instructions. Intracellular staining was performed using antibodies against IL-17A (clone eBio17B7), and RORγt (clone B2D).

Data were acquired on a Thermo Fisher Scientific Attune NxT Flow Cytometer and analyzed using FlowJo software (10.10.0). Gating strategies were established using single-stain controls and are shown in Fig. S1.

##### 16S rRNA gene sequencing

Gnomic DNA was extracted from ileal luminal samples using the Qiagen DNeasy 96 PowerSoil Pro QIAcube HT Kit (cat# 47021). The hypervariable V3-V4 region of the 16S rRNA gene was amplified using primers 341F (5′-CCTAYGGGDBGCWGCAG-3′) and 806R (5′-GACTACNVGGGTMTCTAATCC-3′). Amplicon sequences were analyzed using Qiime2 (2024-5) (5). Denoising was performed using DADA2 with the following parameters: trim-left-f = 20, trim-left-r = 25, trunc-len-f = 300, and trunc-len-r = 276; all other parameters were set to default values (6). A total of 2,479,491 high-quality reads from 732 Amplicon sequence variants (ASVs) were generated from 12 samples. ASVs were assigned taxonomy using a SILVA v138 reference database classifier trained on the V3-V4 region using the same primer sequences (7). Alpha and beta diversity metrics were calculated using the q2-diversity plugin with core-metrics-phylogenetic command with a sampling depth of 169,826 sequences per sample. For correlation analyses, count data were centered log-ratio (CLR) transformed using the compositions::clr() function in R (4.5.1). Differential abundance analysis at the genus level was performed using linear discriminant analysis effect size (LEfSe; v1.1.2) implemented via Bioconda, using default parameters (LDA score > 2.0; $p < 0.05$) (8).

##### Quantitative PCR of SFB

Absolute quantification of 16S gene copy numbers of SFB was quantified according to previously described methods (9, 10). Gnomic DNA was extracted from ileal luminal samples using the ZymoBIOMICS™-96 MagBead DNA Kit (Zymo Research, cat# D4302) according to the manufacturer’s instructions. A synthetic double-stranded DNA fragment (gBlock; Integrated DNA Technologies) was designed based on the mouse SFB 16S rRNA reference sequence (ENA accession X77814) (11). Primer binding sites corresponding to 736F (5′-GACGCTGAGGCATGAGAGCAT-3′) and 844R (5′-GACGGCACGGATTGTTATTCA-3′) were identified, and a continuous 219-bp fragment spanning the qPCR amplicon (~109 bp) with ~55 bp flanking sequence on each side was extracted directly from the reference without modification. The gBlock was synthesized as linear double-stranded DNA and used to generate standard curves for absolute quantification.

Quantitative PCR was performed using SYBR Green with 0.2 µM of each primer in 20 µL reactions. Cycling conditions were: 95°C for 3 min followed by 40 cycles of 95°C for 5 seconds and 60°C for 30 seconds, with melt curve analysis to confirm specificity. Ten-fold serial dilutions of the gBlock were used to generate standard curves ($10^8$-$10^2$ copies per reaction). Copy number was calculated based on fragment length and DNA mass. Absolute SFB abundance was determined by interpolation from the standard curve and normalized to input sample mass.

**Statistical analysis**

Statistical analyses were performed using GraphPad Prism software (v10), QIIME2 (2024.5), and R (4.5.1). For comparisons between two groups, unpaired Welch's t tests were used unless otherwise specified. Microbial alpha diversity metrics were analyzed using the Mann-Whitney U test. Comparisons among three groups were performed using one-way analysis of variance (ANOVA). For 16S rRNA gene analysis, beta diversity differences were assessed using Permutational Multivariate Analysis of Variance (PERMANOVA) and Permutational Analysis of Multivariate Dispersions (PERMDISP) implemented using the q2-diversity plugin with beta-group-significance command in QIIME2. Correlation analyses between centered log-ratio (CLR)-transformed genus-level abundances and Th17 markers were performed using Pearson correlation in R (WGCNA package) (12). Correlations between SFB abundance and either *C. rodentium* burden or Th17 markers were calculated using GraphPad Prism. Survival analysis was performed using the log-rank (Mantel-Cox) test in Graphpad Prism. A $p$-value < 0.05 was considered statistically significant.
